# Parkinson’s LRRK2-G2019S risk gene mutation drives sex-specific behavioral and cellular adaptations to chronic variable stress

**DOI:** 10.1101/2024.06.05.597647

**Authors:** Christopher A. Guevara, Kumayl Alloo, Swati Gupta, Romario Thomas, Pamela del Valle, Alexandra R. Magee, Deanna L. Benson, George W. Huntley

## Abstract

Anxiety is a psychiatric non-motor symptom of Parkinson’s that can appear in the prodromal period, prior to significant loss of brainstem dopamine neurons and motor symptoms. Parkinson’s-related anxiety affects females more than males, despite the greater prevalence of Parkinson’s in males. How stress, anxiety and Parkinson’s are related and the basis for a sex-specific impact of stress in Parkinson’s are not clear. We addressed this using young adult male and female mice carrying a G2019S knockin mutation of leucine-rich repeat kinase 2 (*Lrrk2*^G2019S^) and *Lrrk2*^WT^ control mice. In humans, *LRRK2*^G2019S^ significantly elevates the risk of late-onset Parkinson’s. To assess within-sex differences between *Lrrk2*^G2019S^ and control mice in stress-induced anxiety-like behaviors in young adulthood, we used a within-subject design whereby *Lrrk2*^G2019S^ and *Lrrk2*^WT^ control mice underwent tests of anxiety-like behaviors before (baseline) and following a 28 day (d) variable stress paradigm. There were no differences in behavioral measures between genotypes in males or females at baseline, indicating that the mutation alone does not produce anxiety-like responses. Following chronic stress, male *Lrrk2*^G2019S^ mice were affected similarly to male wildtypes except for novelty-suppressed feeding, where stress had no impact on *Lrrk2*^G2019S^ mice while significantly increasing latency to feed in *Lrrk2*^WT^ control mice. Female *Lrrk2*^G2019S^ mice were impacted by chronic stress similarly to wildtype females across all behavioral measures. Subsequent post-stress analyses compared cFos immunolabeling-based cellular activity patterns across several stress-relevant brain regions. The density of cFos-activated neurons across brain regions in both male and female *Lrrk2*^G2019S^ mice was generally lower compared to stressed *Lrrk2*^WT^ mice, except for the nucleus accumbens of male *Lrrk2*^G2019S^ mice, where cFos-labeled cell density was significantly higher than all other groups. Together, these data suggest that the *Lrrk2*^G2019S^ mutation differentially impacts anxiety-like behavioral responses to chronic stress in males and females that may reflect sex-specific adaptations observed in circuit activation patterns in stress-related brain regions.

## 1 Introduction

Anxiety is a prevalent psychiatric non-motor symptom of genetic and idiopathic forms of Parkinson’s that can appear in the prodromal stage, before significant dopamine neuron degeneration (Shiba et al., 2000; Broen et al., 2016). Chronic behavioral stress has emerged as an important risk factor for psychiatric symptoms and Parkinson’s (Hemmerle et al., 2012). How stress, psychiatric symptoms and Parkinson’s are related mechanistically is unknown. Mice carrying gene mutations associated with an increased risk for Parkinson’s provide an opportunity to examine the intersection between genetic and experience-dependent risks.

The G2019S mutation in *LRRK2* (encoding leucine-rich repeat kinase 2) is among the most common genetic risk mutations for familial and sporadic Parkinson’s (Kluss et al., 2019). *LRRK2*^G2019S^ produces late-onset Parkinson’s that is clinically similar to idiopathic forms, including prodromal anxiety and depression in non-manifesting mutation carriers (Shanker et al., 2011; DeBroff et al., 2023). Genetic, pathological, and preclinical studies suggest that dysregulated LRRK2 may be part of a common pathological mechanism (Kluss et al., 2019; Rocha et al., 2022; Peter and Strober, 2023). In brain, LRRK2 protein is enriched in areas that are susceptible or can adapt negatively to stressful behavioral experiences, including medial prefrontal cortex (mPFC), amygdala, hippocampus and striatum (Simón-Sánchez et al., 2006; Giesert et al., 2013), suggesting that LRRK2 is well-positioned to modify responses to stress.

Previous work has shown that male *Lrrk2*^G2019S^ knockin mice display altered social interaction and hedonic responses following periods of social defeat stress that are distinct from wildtype mice (Matikainen-Ankney et al., 2018; Guevara et al., 2020). Social defeat stress is an intense stressor that has most readily been applied to males. This leads us to ask whether such synergy between social stress and *Lrrk2*^G019S^ can be generalized to different durations and forms of stress. We also address the effects of stress on female *Lrrk2*^G2019S^ mice. To answer these questions, we mounted a fully powered comparison of anxiety-like behaviors and examined brain regions activated in males and females following extended exposure to a chronic variable stress protocol.

## 2 Methods

### Mice

*Lrrk2*^G2019S^ knockin mice (RRID:MGI:6273311) were generated by Eli Lilly and characterized previously (Matikainen-Ankney et al., 2016, 2018; Guevara et al., 2020; Hussein et al., 2022). The knockin mice were congenic on C57Bl/6NTac background, bred as homozygotes, and backcrossed to wildtype C57Bl/6NTac every fifth generation to prevent genetic drift. Age, sex- and strain-matched wildtype mice (*Lrrk2*^WT^) were used as controls. All mice were bred in Mount Sinai’s vivarium under identical conditions. The mice were group housed and maintained on a 12h light/dark cycle with free access to food and water unless otherwise specified. Genotypes of all animals were confirmed through PCR and sequencing. Male and female heterozygous *Lrrk2*^G2019S^ mice and age- and strain-matched wildtype mice were utilized for all experiments. Mice were aged postnatal day 56 - 60 at the start of the behavioral experiments. In the females, no attempt was made to document the stage of the estrous cycle in relation to behavioral testing. Recent studies suggest the impact of the estrous cycle on behavioral performance in tests of anxiety, including those used in this study, is negligible (Chari et al., 2020; Johnson et al., 2021; Levy et al., 2023; Li et al., 2023). All experiments were approved by Mount Sinai’s Institutional Animal Care and Use Committee and conformed to National Institutes of Health guidelines.

### Experimental design of behavioral testing

We used a within-subject design to evaluate the effects of 28d of variable stress on different measures of anxiety-like behaviors. To establish a pre-stress baseline of responses, behaviorally naïve *Lrrk2*^G2019S^ and *Lrrk2*^WT^ mice were tested on a battery of standard tests on days 1 - 3. Twenty-four hours later, mice were subjected to 28d of variable stress. Twenty-four hours after the last day of the stress paradigm, mice were re-tested on the same daily pre-stress behavioral battery to establish post-stress effects for comparison (**Fig. 1**). The pre- and post-stress behavioral assays comprised an open field test, a social interaction test with a same-sex conspecific, and a novelty-suppressed feeding test. One test was conducted per day and animals were allowed to habituate to the testing room for 1h prior to testing. Equipment was cleaned with 70% ethanol in between testing sessions. Mice were group-housed throughout the entire experiment (Westenbroek et al., 2005; Manouze et al., 2019).

**Figure 1.**
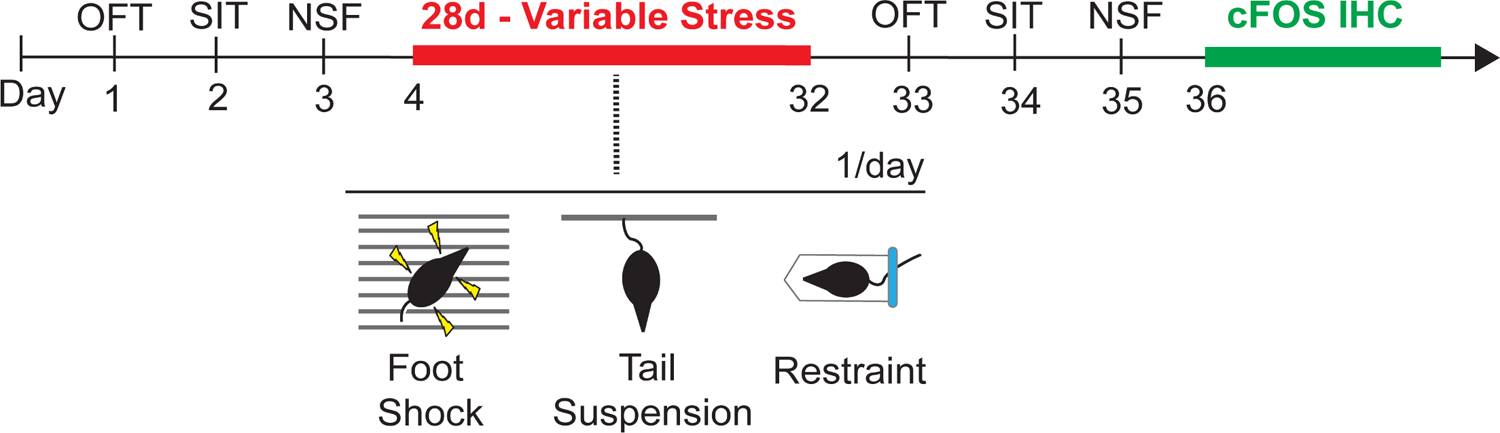
Schematic showing timeline of behavioral tests prior to (baseline) and following a 28d variable stress paradigm, with subsequent cFOS immunohistochemistry (IHC). OFT: Open Field Test; SIT: Social Interaction Test; NSF: Novelty Suppressed Feeding

### Open Field Test (OFT)

Mice were placed in an open arena under white light and allowed to explore for 5 minutes (Cunningham et al., 2021). Fusion Software (v5.0) (Omnitech Electronics) was used to quantify time spent in the periphery and center of the test arena as an indicator of anxiety (Badimon et al., 2020), and to monitor total distance traveled.

### Social Interaction Test (SIT)

Social interaction (SI) was assessed in the absence and presence of a novel conspecific social target under red lights (Muir et al., 2020; Tsyglakova et al., 2021). SI behavior was calculated as an SI Ratio = t_present_/t_absent_, where t_present_ and t_absent_ represent the times spent in the interaction zone in the presence and absence of the novel social target. Time and location in the arena were recorded by video tracking software (Ethovision 10.0, Noldus).

### Novelty Suppressed Feeding (NSF)

NSF tests were conducted as described (Freire-Cobo and Wang, 2020; Johnson et al., 2021). Briefly, mice were food-deprived the night before the test. On the day of testing, mice were placed in an arena containing a single food pellet in the center. The latency to take a bite from the piece of food was scored manually for a maximum of 10 mins. The test was conducted under red light. All mice were weighed in the evening prior to food restriction and the next morning prior to the NSF test.

### Variable stress paradigm

The variable stress paradigm was performed as previously described (Freire-Cobo and Wang, 2020; Johnson et al., 2021). Mice were subjected to one of three stressors in the following order for 1 h per day: 100 random foot shocks at 0.45mA/sec (day 1), tail suspension (day 2), and restraint by placing the animal in a ventilated 50-mL conical tube (day 3). Stressors were administered and repeated in this order over 28d. Mice were returned to their homecage and group-housed at the end of each stressor.

### cFos activity mapping and immunohistochemistry

Twenty-four hours following the last behavioral assay, 3–4 mice of each sex and genotype were subjected to restraint stress for one hour then returned to their homecage for 90 mins followed by transcardial perfusion with 4% paraformaldehyde in phosphate buffered saline (PBS, pH7.3) (**Fig. 1**). Following 6 h postfixation, brains were sectioned coronally at 50 μM on a freezing sliding microtome (Leica). Immunocytochemical localization of cFos was performed on every sixth section (approximately every 300 µm). Free-floating sections were incubated in 2% bovine serum albumin (BSA) in 0.3% Tween-20 PBS for 1 h then incubated in rabbit anti-c-Fos antibody (1:500, Synaptic Systems, catalog # 226-003, RRID: AB_2231974) diluted in the same solution for 72 hours at 4°C. Sections were washed, then incubated in goat anti-rabbit Alexa 647-conjugated antibodies (1:500, Abcam catalog #150076, RRID: AB_2782993) diluted in 2% BSA in 0.3% Tween-20 PBS for 1 h and then stained with 0.005% DAPI in PBS for 20 min. Finally, sections were slide-mounted and coverslipped with Mowiol.

### Data and image analysis

Single-plane, tiled images of entire cFos-immunolabeled brain sections were captured on a Leica DMi8 microscope at 10X magnification. See Results for stress-relevant brain regions analyzed. For each region chosen for analysis (see Results), a region of interest (ROI) was generated in each section using Nissl staining as a guide (Franklin and Paxinos, 2007). ROIs were kept the same size across mice. ROIs in prelimbic (PL) and infralimbic (IL) cortex included layers 1-6; ROIs in hippocampus included all layers.

Density of cFos-labeled cells was quantified using the “Cell Counter” plug-in on ImageJ/Fiji. Thresholding limits were determined using the “Moments” setting on ImageJ/Fiji. cFos density was calculated as:

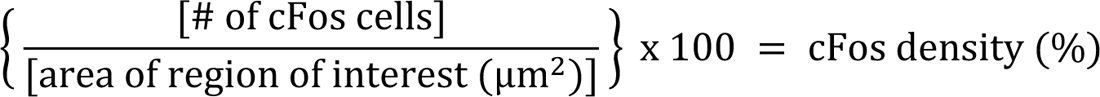

### Statistical analyses

Statistical comparisons of normally distributed data between two groups were conducted using two-tailed Student’s t-tests. To compare parametric data from multiple groups, repeated-measures two-way ANOVA was performed, followed by Šidák’s post-hoc test for multiple comparisons. For nonparametric data between two groups, a Mann–Whitney U test was used. Analyses were conducted using GraphPad Prism (v8.2.1), with p < 0.05 considered significant. Data are presented as means ± SEM.

## 3 Results

We subjected young adult male and female *Lrrk2*^WT^ and heterozygous *Lrrk2*^G2019S^ knockin mice to a variable stress (VS) paradigm for 28 days (28d-VS) to assess the impact of stress, sex and genetic risk for Parkinson’s on measures of anxiety-like behaviors in mice using a within-subject design to compare baseline (pre-stress) and post-stress responses (**Fig. 1**).

### Open Field Test (OFT)

In an OFT, reduced time spent in the center of an open arena is interpreted as increased anxiety-like behavior (Simon et al., 1994; Harro, 2018). At baseline, we found no significant genotype differences among the male and female cohorts in the time spent in the center (**Fig. 2A, D**) or in total distance traveled, indicating that the mutation alone does not significantly affect anxiety-like behavior or locomotor activity in this test in males or females, as expected (Yue et al., 2015). After 28d-VS, male *Lrrk2*^WT^ and *Lrrk2*^G2019S^ mice exhibited no significant stress-related changes in the time spent in the center (**Fig. 2A**). However, in contrast to males, both *Lrrk2*^WT^ and *Lrrk2*^G2019S^ female mice showed a main effect of stress by a significantly decreased time in center following 28d-VS (**Fig. 2D**). There were no significant effects of stress on locomotor activity in males or females (males: main effect of stress, F_1,18_ = 1.267, *p* = 0.054, main effect of genotype, F_1,18_ = 0.1268, *p* = 0.7259; females: main effect of stress, F_1,17_ = 1.807, *p* = 0.1966, main effect of genotype, F_1,17_ = 1.063, *p* = 0.3169).

**Figure 2.**
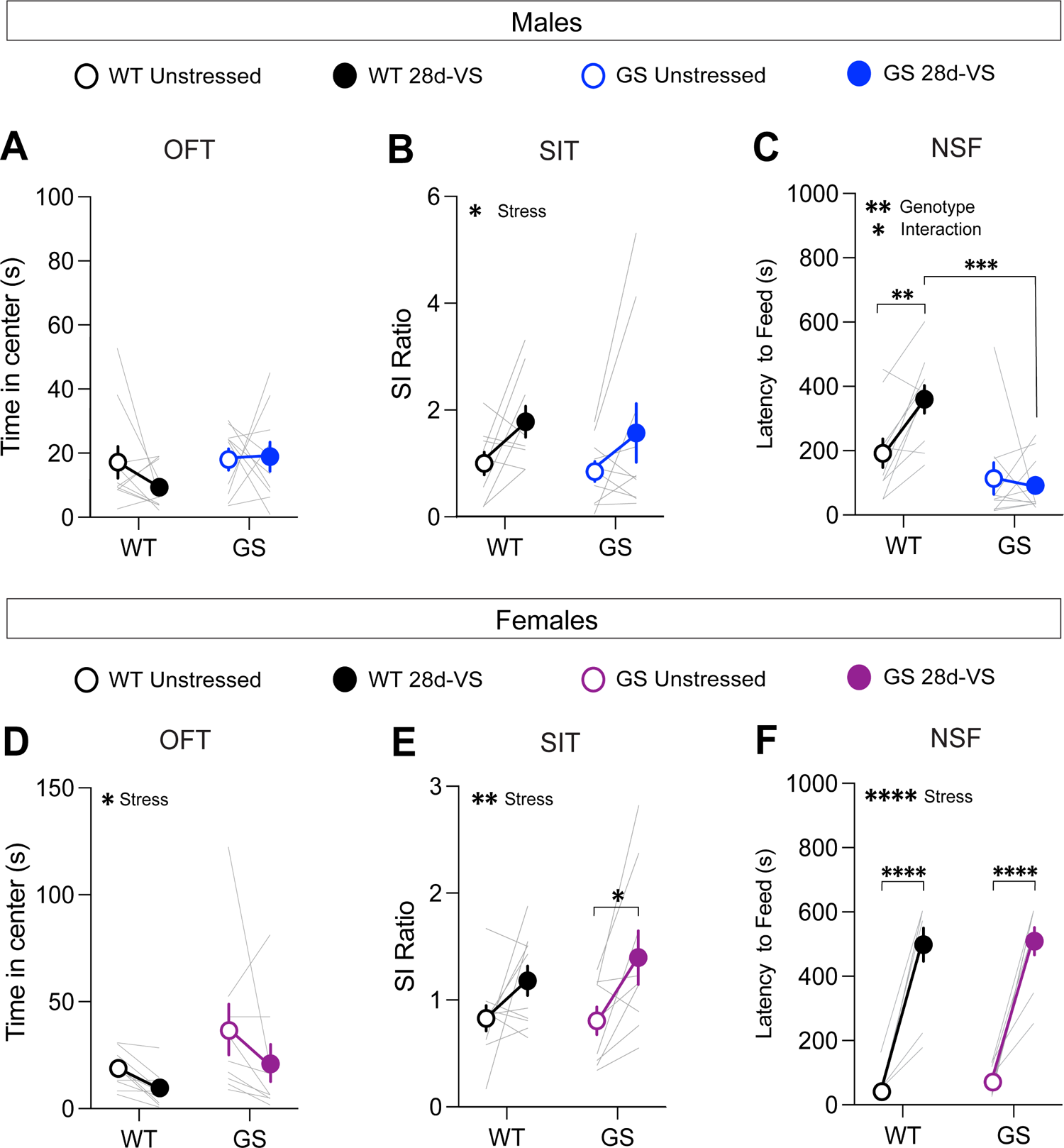
Effects of 28d-VS (variable stress) on measures of anxiety-like behavior, social interaction and hyponeophagia in males (top row) and females (bottom row). (**A**) Male *Lrrk2*^WT^ (WT) control and *Lrrk2*^G2019S^ (GS) mice showed no changes to time in center following 28d-VS in comparison with their baseline measures (*p* = 0.4293, F_1,18_ = 0.6539 for main effect of stress). (**B**) WT and GS males both exhibited an increased social interaction (SI) ratio following 28d-VS in comparison with baseline SI ratios ( *p* = 0.0201, F_1,17_ = 6.572 for main effect of stress). (**C**) In an NSF test, GS males were unaffected by 28d-VS compared to their baseline latencies. In contrast, WT males showed a significantly increased latency to feed (*p* = 0.0099), which was significantly longer than post-stress latencies of GS males (*p* = 0.0001) (*p* = 0.0199, F_1,18_ = 0.2994 for interaction effect; *p* = 0.0014, F_1,18_ = 14.21 for main effect of genotype). (**D**) Both WT and GS females show a decreased time in center following 28d-VS (*p* = 0.046, F_1,17_ = 4.635 for main effect of stress). (**E**) WT and GS females both showed an increased SI ratio following 28d-VS (*p* = 0.0036, F_1,17_ = 11.42 for main effect of stress); with GS females a significantly increased SI ratio compared to their baseline measures (*p* = 0.019). (**F**) Both WT and GS female mice exhibited a similarly increased latency to feed following 28d-VS (*p* = <0.0001) (*p* = <0.0001, F_1,17_ = 143.7 for main effect of stress). n = 10 WT male mice, 10 WT female mice, 13 GS male mice, and 9 GS female mice. Statistical tests used: 2-way repeated measures ANOVA with post hoc Šidák test for multiple comparisons. Data presented as mean ± s.e.m. \**p* < 0.05; \*\**p* < 0.01; \*\*\**p* < 0.001, \*\*\*\**p* < 0.001.

### Social interaction Test (SIT)

Mice were tested for social interaction (SI) behavior with an age- and sex-matched conspecific because social avoidance is a measure of anxiety- and depression-like states (Golden et al., 2011). There were no baseline genotype differences in SI ratios in males or females (**Fig. 2B, E**), as expected (Matikainen-Ankney et al., 2018; Guevara et al., 2020). However, there was a main effect of 28d-VS on SI behavior in both genotypes and sexes (**Fig. 2B, E**). In males and females, we found that 28d-VS significantly *increased* social interaction in *Lrrk2*^WT^ and *Lrrk2*^G2019S^ mice. Posthoc analysis of the female cohort showed that female *Lrrk2*^G2019S^ mice had a significantly greater increase in SI ratios following 28d-VS compared to their baselines (**Fig. 2E**).

### Novelty Suppressed Feeding (NSF)

Mice were food-restricted overnight and subsequently placed in a novel arena. Their latency to feed on a new food pellet was measured. Increased latency to feed is interpreted as a measure of heightened anxiety-like behavior because this measure is reduced by anxiolytic drugs (Bodnoff et al., 1989). Under baseline conditions, there were no genotype differences in males or females in response latencies to feed (**Fig. 2C, F**). In males, posthoc analyses showed that following 28d-VS, *Lrrk2*^WT^ mice exhibited significantly increased latencies to feed compared to their pre-stress baseline responses (**Fig. 2C**). In contrast, male *Lrrk2*^G2019S^ showed no significant differences between unstressed and post-stress latencies to feed (**Fig 2C**), suggesting a lack of anxiety-like effect in this test driven by 28d-VS compared to a significant elevation in anxiety-like behavior of *Lrrk2*^WT^ mice. We verified that in the male mice, there were no genotype differences in the percent body weight lost after overnight fasting, either under baseline or post-28d-VS conditions (repeated-measures two-way ANOVA, main effect of genotype, F_1,18_ = 0.2795, *p* = 0.6035), ruling out potential differences between genotypes in weight lost as a factor accounting for different post-stress effects on latency to feed.

In female mice, there was a significant effect of stress on latencies to feed that was evident in both genotypes, with *Lrrk2*^WT^ and *Lrrk2*^G2019S^ female mice exhibiting significantly increased latencies to feed following 28d-VS compared to their unstressed baseline latencies (**Fig. 2F**).

Comparison of overnight fasting-induced body weight loss showed that at baseline, female *Lrrk2*^G2019S^ mice lost a significantly greater percentage of body weight compared to baseline measured in female wildtype mice (two-way RM ANOVA, main effect of genotype, F_1,17_ = 9.262, *p* = 0.0073), but there were no significant differences between genotypes in body weight lost following 28d-VS. As the latencies to feed were similar between genotypes under both baseline and post-stress conditions, the significance of a greater percentage of body weight lost at baseline in the *Lrrk2*^G2019S^ females is not clear. In sum, following stress, latency to feed was increased significantly in both male and female wildtype mice and in female *Lrrk2*^G2019S^ mice. Male *Lrrk2*^G2019S^ mice were unaffected. A summary of behavioral results, following 28d-VS in males and females, is shown in **Fig. 3**.

**Figure 3.**
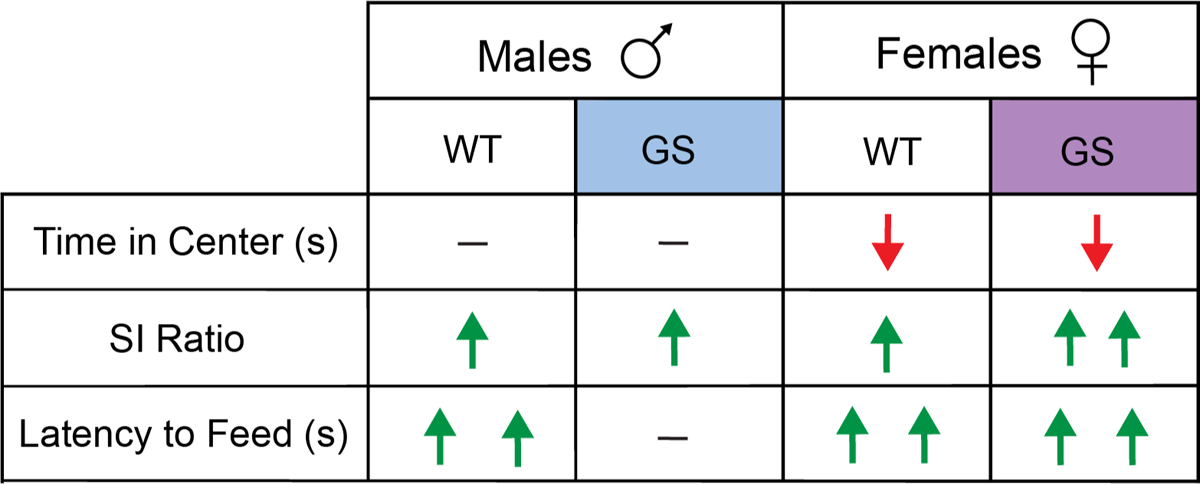
Summary of effects of 28d-VS on WT and GS male and female mice across all behavioral measures. Directionality of change in behavioral performance following 28d-VS is indicated by color and direction of arrows. Red downward arrow = decreased measure of behavior performance following 28d-VS compared to baseline measures; green upward arrow = increased measure of behavioral performance following 28d-VS compared to baseline measures; horizontal line = no change in measure of behavioral performance following 28d-VS. More than one arrow = significance in post-hoc comparison.

### Cellular activity mapping by cFos immunolabeling following 28d-VS

Brain regions implicated in anxiety-like behaviors following chronic stress in humans and mice include mPFC (areas PL and IL in mice), dorsal striatum (dStr), nucleus accumbens (NAc), basolateral amygdala (BLA), dorsal hippocampal area CA1 (dHPC) and ventral hippocampal area CA1 (vHPC). To assay sex and genetic differences in regional activation patterns of the stress-network following 28d-VS, we used immunolabeling for the immediate early gene cFos as a neural activity marker (Morgan and Curran, 1989) (**Fig. 4A**).

**Figure 4.**
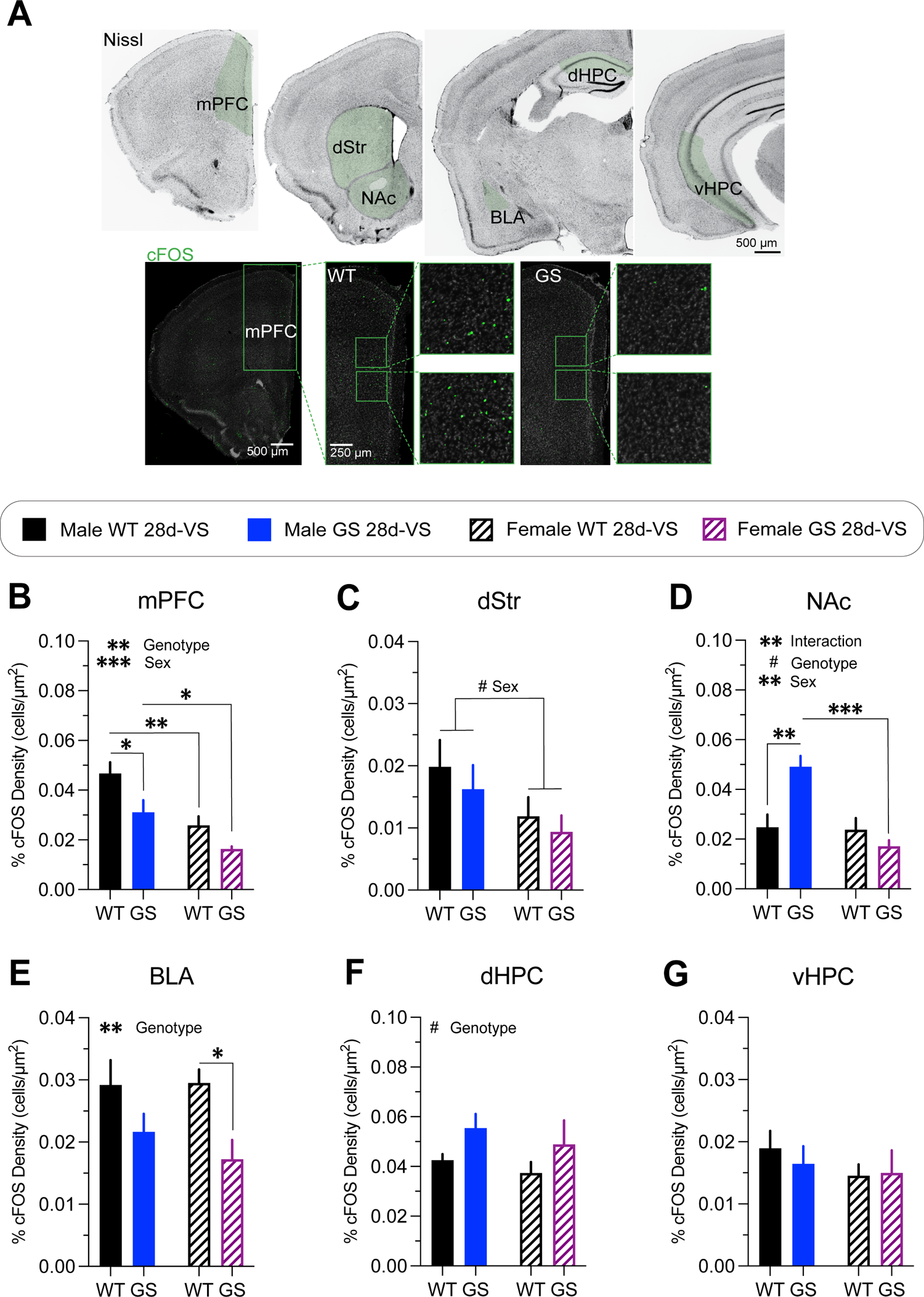
cFos immunolabeling-based cellular activity patterns in male and female mice following 28d-VS. cFos immunofluorescent labeling of perfused brain sections was used as an activity marker in stress-relevant brain regions following 28d-VS. (**A**) A Nissl-stained series of frontal sections (top row) through the brain of a wildtype mouse post-stress with regions in green shading highlighting those analyzed for cFos immunolabeling. Representative confocal microscope images through mPFC of a male WT mouse (bottom row, left) or of a male GS mouse (bottom row, right) showing examples of cFos immunofluorescently-labeled cells (boxed regions show higher-magnification images). mPFC, medial prefrontal cortex; dStr, dorsal striatum; NAc, nucelus accumbens; BLA, basal amygdala; dHPC, dorsal hippocampus; vHPC, ventral hippocampus. Following 28d-VS, mice exhibited differential genotype and sex differences in most areas examined. (**B**) Analysis of cFos-labeled cell density in mPFC post-stress (*p* = 0.0085, F_1,14_ = 9.345 for genotype effect; *p* = 0.0006, F_1,14_ = 19.06 for main effect of sex); with GS males (*p* = 0.0258) and WT females (*p* = 0.0057) showing decreased density of cFos-labeled cells compared to WT males, and GS females showing decreased density of cFos-labeled cells compared to GS males (*p* = 0.0442). (**C**) Analysis of cFos-labeled cell density in dStr post-stress (*p* = 0.0657, F_1,14_ = 3.985 for main effect of sex). (**D**) Analysis of cFos-labeled cell density in NAc post-stress (*p* = 0.0037, F_1,14_ = 12.05 for interaction effect; *p* = 0.0672, F_1,14_ = 3.936 for genotype effect; *p* = 0.0024, F_1,14_ = 13.68 for main effect of sex); with GS males (*p* = 0.0022) showing increase cFos-labeled cell density compared to WT males, and GS females showing decreased density of cFos-labeled cells compared to GS males (*p* = 0.0003). (**E**) Analysis of cFos-labeled cell density in BLA post-stress (*p* = 0.0088, F_1,14_ = 9.261 for genotype effect); with GS females showing a significantly decreased density of cFos-labeled cells compared to WT females (*p* = 0.0474). (**F**) Analysis of cFos-labeled cell density in dHPC post-stress (*p* = 0.0667, F_1,13_ = 4.005 for main effect of genotype). (**G**) Analysis of cFos-labeled cell density in vHPC post-stress mice (*p* = 0.62, F_(1,14)_ = 0.25 for interaction effect; *p* = 0.72, F_(1,14)_ = 0.13 for genotype effect; *p* = 0.33, F_(1,14)_ = 0.10 for sex effect). For all experiments, n = average of 2 brain slices per mouse / 3-4 mice per group. Statistical tests used: Statistical tests used: 2-way repeated measures ANOVA with post hoc Šidák test for multiple comparisons. Data presented as mean ± s.e.m. ^#^*p* = 0.05-0.07, \**p* < 0.05; \*\**p* < 0.01; \*\*\**p* < 0.001, \*\*\*\**p* < 0.001.

Both male and female *Lrrk2*^G2019S^ mice had a significantly decreased density of cFos immunofluorescent cells in mPFC following 28d-VS compared to their respective 28d-VS *Lrrk2*^WT^ controls (**Fig. 4B**), with combined males having a significantly higher cFos-labeled cell density overall in comparison with combined females (**Fig. 4B**). In dStr (**Fig. 4C**), there was a trending main effect of sex with combined males having higher densities of cFos-labeled cells compared to combined females. In NAc (**Fig. 4D**), there was a significant increase in the density of cFos immunolabeled cells in male *Lrrk2*^G2019S^ mice compared to male *Lrrk2*^WT^ mice, but no significant differences in cFos-labeled cell density between *Lrrk2*^G2019S^ and *Lrrk2*^WT^ female mice (**Fig. 4D**).

Together, these data support a main effect of genotype, sex and their interaction. In BLA (**Fig. 4E**), there was a main effect of genotype, with lower cFos-labeled cell densities in *Lrrk2*^G2019S^ mice compared to *Lrrk2*^WT^ mice and significantly lower cFos-labeled cell density in stressed *Lrrk2*^G2019S^ female mice compared to stressed female *Lrrk2*^WT^ mice. In dHPC (**Fig. 4F**), there was a trending main effect of genotype with higher densities of cFos-labeled cells in *Lrrk2*^G2019S^ mice compared to *Lrrk2*^WT^ controls, while in vHPC, there were no genotype or sex-related differences in cFos-labeled cells following 28d-VS (**Fig. 4G**). Together, these data suggest that the recruitment of neural networks across some stress-responsive brain regions differs between male and female mice. c-Fos-based activation levels appeared to be generally higher in male than in female mice. Most striking was the selectively increased activation of NAc in male *Lrrk2*^G2019S^ mice because, outside of this, the mutation appeared to mostly dampen activation.

## 4 Discussion

The purpose of the study was to better understand how the Parkinson’s *Lrrk2*^G2019S^ risk mutation impacted stress-induced anxiety-like behaviors in young adulthood in males and females. We subjected male and female *Lrrk2*^G2019S^ heterozygous knockin and *Lrrk2*^WT^ control mice to a 28d chronic variable stress paradigm to assess within-sex differences between *Lrrk2*^G2019S^ mice and *Lrrk2*^WT^ controls. Using several tests of anxiety-like behaviors, we found that in males, chronic variable stress affected behavioral performance of *Lrrk2*^WT^ and *Lrrk2*^G2019S^ mice similarly with the exception of a NSF test, where stress had no impact on *Lrrk2*^G2019S^ while significantly affecting wildtypes. Female *Lrrk2*^G2019S^ mice were impacted by chronic stress similarly to *Lrrk2*^WT^ control females across all behavioral measures. Subsequent post-stress analyses compared cFos immunolabeling-based cellular activity patterns across several stress-relevant brain regions. We found that in male *Lrrk2*^G2019S^ mice, post-stress patterns of cFos-labeled cells in several areas, including mPFC and the NAc, differed significantly from those found in either stressed *Lrrk2*^WT^ males or stressed *Lrrk2*^G2019S^ females. In contrast, stressed female *Lrrk2*^G2019S^ mice showed cFos-labeling patterns that were mostly similar to stressed female wildtypes with the exception of the BLA, which had significantly fewer cFos-labeled neurons in stressed female *Lrrk2*^G2019S^ mice compared to stressed female *Lrrk2*^WT^ mice. Together, these data suggest that the *Lrrk2*^G2019S^ mutation differentially impacts anxiety-like behavioral responses to chronic stress in male and female mutation carriers. This may reflect sex-specific adaptations in circuit activation patterns in many, but not all, stress-related brain regions. More broadly, our findings suggest a potential basis for differences in the onset and/or severity of psychiatric symptoms in male and female Parkinson’s patients.

The effects of 28d-VS on anxiety-like behaviors of our *Lrrk2*^WT^ control mice were mostly similar to previous studies of wildtype mice despite some differences in protocols. For example, we found a main effect of stress in the significantly longer latencies to feed in the NSF test in male and female *Lrrk2*^WT^ mice as reported previously with shorter (Bittar et al., 2021) or identical periods of exposure to similar stressors (Johnson et al., 2021). Similarly, social interaction time with a sex-matched, conspecific target increased in both male and female *Lrrk2*^WT^ mice following 28d-VS as reported previously (Zhao et al., 2020; LeClair et al., 2021; Li et al., 2023). Importantly, in the absence of stress, we found no within-sex differences between *Lrrk2*^G2019S^ knockin mice and their wildtype controls at baseline. This observation indicates that the mutation does not, in and of itself, drive aberrant anxiety-like, depression-like or anhedonia-like behaviors in either male or female *Lrrk2*^G2019S^ mice at the young adult ages examined, as suggested previously (Yue et al., 2015; Volta et al., 2017; Matikainen-Ankney et al., 2018; Guevara et al., 2020). While there were no baseline behavioral effects of the mutation at young adult ages, it is possible that in the absence of stress, mutation-driven effects on anxiety- or depression-like behaviors would emerge with advancing age. In a study using BAC transgenic mice overexpressing human *LRRK2*^G2019S^, anxiety- and depression-like behaviors that were absent in young adulthood first appeared in both male and female mutant mice at about 1 year of age—in the absence of overt stress—and persisted at older ages (Lim et al., 2018). Moreover, such age-dependent effects on psychiatric-like behaviors in these transgenic mice were correlated with contemporaneous changes in hippocampal serotonin levels and altered serotonergic receptor expression (Lim et al., 2018). Setting aside significant differences in the genetics of the two mouse models, it may be that the knockin mice may gradually show an age-dependent effect on anxiety-like behaviors in the absence of stress. Future studies will be needed to examine this and to see if serotonergic innervation becomes compromised with advancing age in the knockin mice. In middle-aged human *LRRK2*^G2019^ carriers, alterations in serotonin transporter binding have been reported (Wile et al., 2017).

In idiopathic Parkinson’s, symptoms of anxiety and depression are more prevalent in women than in men (Leentjens et al., 2011; Cui et al., 2017). Although the prevalence of anxiety and depression in human *LRRK2* mutation carriers is similar to that in idiopathic Parkinson’s (Gaig et al., 2014; San Luciano et al., 2017; Chen et al., 2020b; DeBroff et al., 2023), it is less clear whether there are sex differences in psychiatric symptoms among manifesting *LRRK2*^G2019S^ carriers. One study suggests that depression severity may be similar in male and female mutation carriers (San Luciano et al., 2017), which aligns with the equal penetrance of the mutation in male and female carriers (Marder et al., 2015). Our data in the young adult *Lrrk2*^G2019S^ mice suggest that behavioral stress impacts anxiety-like behaviors in females more significantly than in males, bearing some similarity to idiopathic Parkinson’s. Elevated LRRK2 kinase activity, in the absence of genetic mutation, is found in sporadic Parkinson’s (Kluss et al., 2019; Rocha et al., 2022). As such, behavioral outcomes and regional cell activation analyses of stress effects in *Lrrk2*^G2019S^ mice may have broad relevance to Parkinson’s generally.

Despite broadly similar anxiety-like behavioral responses to 28-VS between *Lrrk2*^G2019S^ and wildtype mice, the densities of cFos-labeled cells across brain regions activated under such conditions showed both genotype-related and sex-specific differences. In stressed male and female *Lrrk2*^G2019S^ mice, densities of cFos-labeled cells were affected in mostly similar ways compared to stressed *Lrrk2*^WT^ mice—lower (mPFC, dStr and BLA), slightly higher (dHPC) or unchanged (vHPC). The exception to this pattern was in the NAc. In this structure, cFos densities in male *Lrrk*^G2019S^ mice were significantly higher than all other groups, whereas in female *Lrrk*^G2019S^ mice, cFos density was similar to male and female *Lrrk2*^WT^ mice. It is possible that in male *Lrrk*^G2019S^ mice, the unique combination of dysregulated top-down control of NAc circuitry by mPFC (Britt et al., 2012) and/or the BLA (Stuber et al., 2011) suggested by lowered cFos densities in those structures, coupled with heightened activation of NAc neurons suggested by the elevated density of cFos-labeled cells, reflects abnormal network activity that ultimately increases motivation to feed and/or blunts anxiety associated with the novel arena. Previous studies of wildtype mice have shown that NSF produces elevated densities of cFos-labeled cells in NAc (in comparison with controls exploring an open-field arena), while optogenetically activating an anterior paraventricular thalamus-to-NAc projection during the NSF test increases motivation to consume food in a novel environment and increases time spent in the open arms of an elevated plus maze (Cheng et al., 2018), commonly interpreted as lowered anxiety. Additionally, other forms of chronic stress in wildtype mice increase glutamate release from mPFC terminals onto a subpopulation of BLA neurons that is correlated with heightened anxiety-like behaviors. Decreasing such stress-induced glutamate release reduces anxiety-like behaviors (Liu et al., 2020).

The mechanistic basis for genotype- and sex-specific differences in anxiety-induced behavioral measures and cellular activation patterns remain speculative. It is known that at baseline (in the absence of any overt behavioral challenge), excitatory synapses in striatum of young adult male and female *Lrrk2*^G2019S^ mice do not express LTP and exhibit significant alterations in synaptic glutamate receptor subunit composition and levels compared to wildtype synapses that reflect impaired GluA subunit trafficking (Matikainen-Ankney et al., 2018; Chen et al., 2020a; Gupta et al., 2024). Previous studies have shown that striatal projection neurons in the NAc of male wildtype mice, when exposed to a 1-day social defeat stress protocol, exhibit significant changes in intrinsic excitability but no changes in properties of spontaneous excitatory synaptic transmission.

Conversely, striatal projection neurons in the NAc of male *Lrrk2*^G2019S^ mice exposed to the same social defeat stress protocol exhibit significant changes in properties of spontaneous excitatory synaptic transmission but no changes in intrinsic excitability (Guevara et al., 2020). Such genotype-specific differences in circuit-level responses to a brief period of social stress suggest that the *Lrrk2*^G2019S^ mutation drives different cellular and synaptic adaptations to other types or durations of behavioral stress. While we did not examine cellular or synaptic properties electrophysiologically or neurochemically in the present study, a previous study of wildtype mice exposed to sub-chronic (6d) variable stress using the same stressors showed significant changes in molecular markers of excitatory synapses in the NAc of female, but not male mice (Brancato et al., 2017). Thus, it is likely that 28d-VS drives both genotype- and sex-specific differences in synaptic and non-synaptic adaptations in several, if not all of the stress-relevant brain regions.

Finally, *Lrrk2*^G2019S^ may be differentially influencing stress-relevant brain circuits through exogenous inflammatory mediators. Cells of the peripheral innate immune system express high levels of LRRK2 (Ahmadi Rastegar and Dzamko, 2020), many of which gain access to the brain following periods of behavioral stress, where they play significant roles in modulating cellular and synaptic function in stress-relevant brain regions (Hodes et al., 2015; Weber et al., 2017). Additionally, recent studies show that wildtype mice exposed to a similar 28d-VS protocol display sex-specific disruptions in gut microbiota (Kropp et al., 2024) and intestinal barrier (tight junction) homeostasis (Doney et al., 2024). This may not only underlie genotype- and sex-specific differences in stress-driven access of inflammatory mediators to the bloodstream and subsequently the brain, which is relevant for Parkinson’s, but may also be relevant to LRRK2 mutation-associated risk of developing inflammatory bowel disease (Kars et al., 2024).

## 5 Author Contributions

CAG, KA, SG, PdV, DLB and GWH Designed Research; CAG, KA, SG, RT, PdV and ARM Performed Research; CAG, KA, SG, PdV, DLB and GWH Analyzed Data; CAG, DLB and GWH wrote the paper.

## Acknowledgments

We thank Kyomi Blake, Jamal Magoti, and Amelia Wieland for assistance in behavioral experiments.

